# Bayesian analyses of Yemeni mitochondrial genomes suggest multiple migration events with Africa and Western Eurasia

**DOI:** 10.1101/010629

**Authors:** Deven N. Vyas, Andrew Kitchen, Aida T. Miró-Herrans, Laurel N. Pearson, Ali Al-Meeri, Connie J. Mulligan

## Abstract

Anatomically modern humans (AMHs) left Africa ˜60,000 years ago, marking the first of multiple dispersal events by AMH between Africa and the Arabian Peninsula. The southern dispersal route (SDR) out of Africa (OOA) posits that early AMHs crossed the Bab el-Mandeb strait from the Horn of Africa into what is now Yemen and followed the coast of the Indian Ocean into eastern Eurasia. If AMHs followed the SDR and left modern descendants *in situ*, Yemeni populations should retain old autochthonous mitogenome lineages. Alternatively, if AMHs did not follow the SDR or did not leave modern descendants in the region, only young autochthonous lineages will remain as evidence of more recent dispersals. We sequenced 113 whole mitogenomes from multiple Yemeni regions with a focus on haplogroups M, N, and L3(xM,N) as they are considered markers of the initial OOA migrations. We performed Bayesian evolutionary analyses to generate time-measured phylogenies calibrated by Neanderthal and Denisovan mitogenome sequences in order to determine the age of Yemeni-specific clades in our dataset. Our results indicate that the M1, N1, and L3(xM,N) sequences in Yemen are the product of recent migration from Africa and western Eurasia. Although these data suggest that modern Yemeni mitogenomes are not markers of the original OOA migrants, we hypothesize that recent population dynamics may obscure any genetic signature of an ancient SDR migration.

## INTRODUCTION

Mitochondrial DNA (mtDNA) has been widely used to expand our understanding of the temporal and geographic structure of global human migrations from the time of our most recent maternal common ancestor, roughly 150 to 200 thousand years ago (kya) (Cann et al. 1987; Underhill and Kivisild 2007) to the present. Phylogenetic analyses of mtDNA data allow us to estimate the order and timing of newly evolved mitochondrial lineages and attempt to correlate their evolution with population movements. Notably, the most basal branches of the human mitochondrial tree (e.g., lineages L0 and L1) are entirely of African origin, suggesting that the most recent common ancestor (MRCA) of all human mitochondrial lineages lived in Africa and that the first successful migrations out of Africa (i.e., those leaving modern descendants outside of Africa) occurred around 60 kya. Evidence for the antiquity of the first, successful out of Africa (OOA) migrations come from the estimated time to the MRCA (TMRCA) of mitochondrial macrohaplogroups M and N, which are thought to have evolved soon after migration out of Africa (Quintana-Murci et al. 1999; Kivisild et al. 2004; Underhill and Kivisild 2007). Importantly, these two groups (M and N) are subgroups of a larger East African group known as L3 and are especially interesting as all non-African populations without recent African admixture carry sub-lineages of only groups M and N. The mtDNA-based timing for the first migration out of Africa is supported by analyses of nuclear DNA data also suggesting that the initial diversification of non-Africans occurred around 62-75 kya based on data from the genome sequence of an Australian Aborigine (Rasmussen et al. 2011) as well as estimates that the introgression of Neanderthal alleles into the ancestors of all non-Africans occurred around 47-65 kya (Sankararaman et al. 2012).

In addition to the timing of the initial OOA migration, the route taken by these early travelers is of central importance to our understanding of AMH dispersals. Two hypothesized routes are the Northern Dispersal Route (NDR) and the Southern Dispersal Route (SDR). The NDR hypothesis posits that anatomically modern humans (AMHs) followed the Nile into what is now Egypt, and entered the Levant near the northern extent of the Red Sea (Quintana-Murci et al. 1999; Macaulay et al. 2005; Rowold et al. 2007). On the other hand, the SDR hypothesis posits that AMHs left Africa through the Horn of Africa by crossing the Red Sea at the Strait of Bab el-Mandeb and dispersing along the coast of Indian Ocean (Quintana-Murci et al. 1999; Metspalu et al. 2004; Macaulay et al. 2005; Thangaraj et al. 2005; Rowold et al. 2007). Ancient, basal clades from macrohaplogroups M and N that predate 50 kya have been found throughout South and Southeast Asia and have been used as evidence in support of the SDR (Endicott et al. 2003; Macaulay et al. 2005; Thangaraj et al. 2005; Oppenheimer 2012). Interestingly, no M or N lineages with very ancient dates have thus far been found in southern Arabia, which would have been the first step in the SDR. The sequences that are currently available from southern Arabia are from more recently evolved haplogroups such as R0a and HV, and thus are more informative for recent migrations than ancient migrations (Kivisild et al. 2004; Černý et al. 2008; Černý et al. 2011; Musilová et al. 2011; Al-Abri et al. 2012). To shed light on the NDR vs. SDR debate, fully characterizing the mitochondrial genetic diversity of southern Arabian populations is critical to determine if they are descendants of an ancient OOA migration, as well as investigating subsequent dispersals in the region.

In this study, we sequenced 113 Yemeni whole mitochondrial genomes (henceforth referred to as mitogenomes) from a set of 550 samples collected across Yemen in 2007. Our strategy was to sequence all Yemeni samples that could carry ancient mitochondrial haplogroups based on initial hypervariable region I (HVR I) classification. Thus, macrohaplogroup M and N samples were chosen for whole mitogenome sequencing as well as all samples from other branches of macrohaplogroup L3 [henceforth referred to as L3(xM,N)]. Some subgroups of M and N, such as D, I, J, and R0a, were excluded since they are more derived (and thus younger) lineages (van Oven and Kayser 2009). Notably, our study represents the most comprehensive analysis of whole mitogenome data from the ancient mitochondrial haplogroups in southern Arabia. We combine our 113 mitogenomes with 338 previously published mitogenomes from surrounding regions and beyond to provide the most comprehensive test yet of the SDR based on local mitochondrial variation. Specifically, we explore whether subsets of Yemeni, Horn of Africa, and Arabian mitogenomes cluster in monophyletic clades that exhibit ancient divergence dates, which are predicted given an ancient dispersal along the SDR. Critically, we use a Bayesian coalescent framework with a relaxed clock and data partitioning (Drummond et al. 2006; Drummond et al. 2012) as opposed to the more commonly used ρ statistic (Forster et al. 1996; Soares et al. 2009), which has been argued to possibly produce inaccurate divergence date estimates (Cox 2008). We used whole mitogenome sequences from Neanderthal and Denisovan remains to provide a more informative temporal calibration for our molecular clock (Green et al. 2008; Briggs et al. 2009; Krause et al. 2010; Reich et al. 2010). Furthermore, unlike many Bayesian coalescent studies of human mitochondrial data, we utilized all of the information contained in the mitogenome sequences, not just the coding region (Atkinson et al. 2008; Gunnarsdóttir et al. 2011). Finally, we interpret the ages and distribution of Yemeni mitogenome lineages to test for support of an SDR migration, as well as to investigate other later dispersals.

## RESULTS

### Mitogenome sequences – results of Sanger and indexed library sequencing

Sanger sequencing of the initial 23 samples yielded 14 complete mitogenome sequences, 8 nearly complete mitogenome sequences with an average of 25 un-sequenced base pairs, and one mostly complete mitogenome missing ˜ 350 bps of sequence (Y502). For the indexed library sequencing, a single flow cell yielded a total of 165,740,000 reads of which 161,400,000 were correctly tagged and 4,340,000 reads were untagged. After trimming adapter sequences and joining paired-ends, each of the 90 Yemeni samples had on average about 2.3 million reads. For each sample, an average of 55.57% of reads (with a minimum of 40.65% and a maximum of 73.06%) were successfully aligned to the rCRS, indicating a successful enrichment procedure of approximately one order of magnitude, as the mtDNA to nDNA ratio in normal human saliva is thought to be around 0.0537:1 (Kim et al. 2004). The number of reads aligned per sample is depicted in Figure S1. The mean coverage for each sample was between 1,204× and 21,253×. The lowest coverage at any given base was 253×. Minimum, maximum and mean coverage per sample is listed in Table S1.

### Mitogenome sequences – haplogroup classification

The 113 sequenced samples yielded 106 different mitogenomes, which were classified into 57 specific haplogroups (Table S2). From the 113 samples, 100 yielded haplogroups consistent with classifications from HaploGrep (Kloss-Brandstätter et al. 2011) based on the existing HVR I sequence, and thirteen yielded haplogroups that were significantly different (e.g., the HVR I sequence indicated haplogroup M43b for sample Y247, but the mitogenome sequence indicated H15). Discordant classifications persisted even when the original HVR I sequences were re-classified using MitoTool (Fan and Yao 2013). For eleven of these thirteen mitogenomes, the HVR I sequences that we generated were completely identical to the existing sequences at all positions (with the exception of the phylogenetically uninformative site 16182) (van Oven and Kayser 2009). Our data suggest that HVR I sequences may be insufficient for accurate mitochondrial haplogroup classification in more than 10% of cases.

From the 113 samples, a total of 106 different sequences were generated in this study; 11 sequences (n=12 samples) were classified as haplogroup L0, 37 sequences (n=39) as L3(xM,N), 20 sequences (n=21) as M1, 14 sequences (n=16) as N1, and 24 sequences (n=25) were classified to other haplogroups. A total of thirteen subhaplogroups were identified three or more times: M1a1 (n=8 samples), L0a2a2a (n=7), N1a1a3 (n=6), L3b1a1a (n=5), L3d1a1a (n=5), L3x1 (n=5), M1a1b (n=5), L3h2 (n=4), L3i2 (n=4), M1a5 (n=4), L0a2c (n=3), N1a3 (n=3), and N1b1a (n=3). A total of 44 haplogroups were represented by only one or two samples; these samples come from a wide array of haplogroups, but due to their status as singletons and doubletons, they were minimally informative in our analyses since our goal was to look for monophyly of Yemeni samples. It is important to note that 20 of the singleton/doubleton samples came from haplogroups not targeted in this study, and thus may not be singletons/doubletons in our overall Yemeni sample set. Seven pairs of Yemeni sequences were identical to each other, classified to L0a2a2a, L3b1a1a, L3d1a1a, M1a1b, M52b, N1a3, and N1b1a. In four cases the pairs were from the same sampling sites, in one case the pair was from the same Yemeni governate but different sampling sites, and in two cases the pairs were from different governates. Additionally, four of our Yemeni sequences were found to be identical to previously published sequences from the United States, Somalia, and Yemen (Socotra) in haplogroups L3b1a, L3d1a1a, and L3h2 (n=2) (Just et al. 2008; Soares et al. 2012). Consistent with previous findings by Černý et al. (2008) who also used this collection of samples, five sequences (n=6) were classified to South Asian haplogroups M3a2, M5a2a2, M5a5, M44a1, and M52b (n=2).

### Bayesian Phylogenetic Analysis of Yemeni Sequences

All results reported below derive from the analysis that excluded the additional non-M1/N1 sequences and included the Denisovan sequences (n=344; see Materials and Methods). The full phylogeny is shown in Figure 1 and detailed phylogenies of haplogroups L0, L3(xM,N), M1, and N1 are shown in Figure 2a-d, respectively. Color blocks are used to delineate the featured subhaplogroups and are the same in all phylogenies. An annotated phylogeny with dates and posterior probabilities for all nodes is shown in Figure S2, and an annotated phylogeny with height 95% HPD bars for all nodes) is shown in Figure S3.

**Figure 1.**
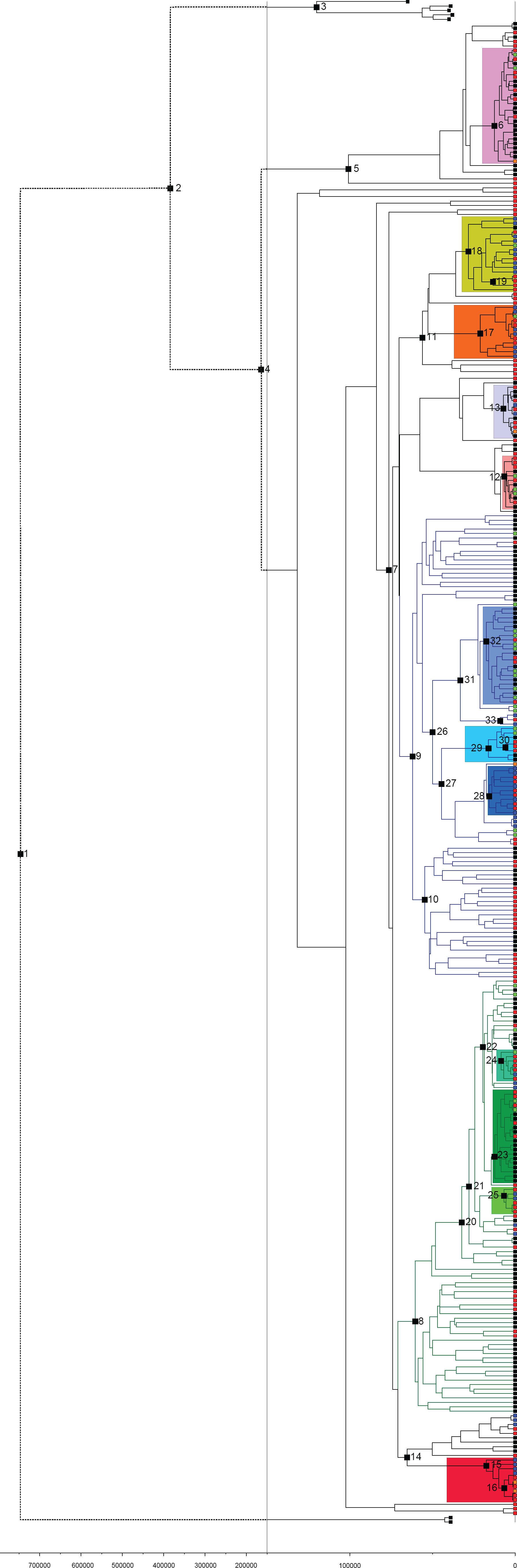
The entire phylogeny from the Bayesian analysis is depicted. All nodes numbers correspond to the numbers listed in Table 1. The time scale and branches greater than 150kya are compressed relative to the rest of the phylogeny and are represented with dashed lines. All branches of macrohaplogroup M are colored dark green and all branches of macrohaplogroup N are colored dark blue. Blocks of color are used to highlight the twelve well-represented (n≥3) subhaplogroups that are analyzed in this study and shown in detail in Figure 2a-d; shades of green highlight closely related M1 groups, shades of blue highlight closely related N1 groups, and different colors highlight the distantly-related L0 and L3(xM,N) groups. Branch tips are colored to indicate the source of each sequence as follows: Red = Yemeni samples from the sample set of C.J.M.; Orange = Yemeni samples from GenBank; Blue = East African samples (i.e., samples from the Horn of Africa and immediately neighboring countries); Green = samples from the Near East (i.e., samples from the Arabian Peninsula, the Levant, and Turkey); and Black = all other regions. .

**Figure 2.**
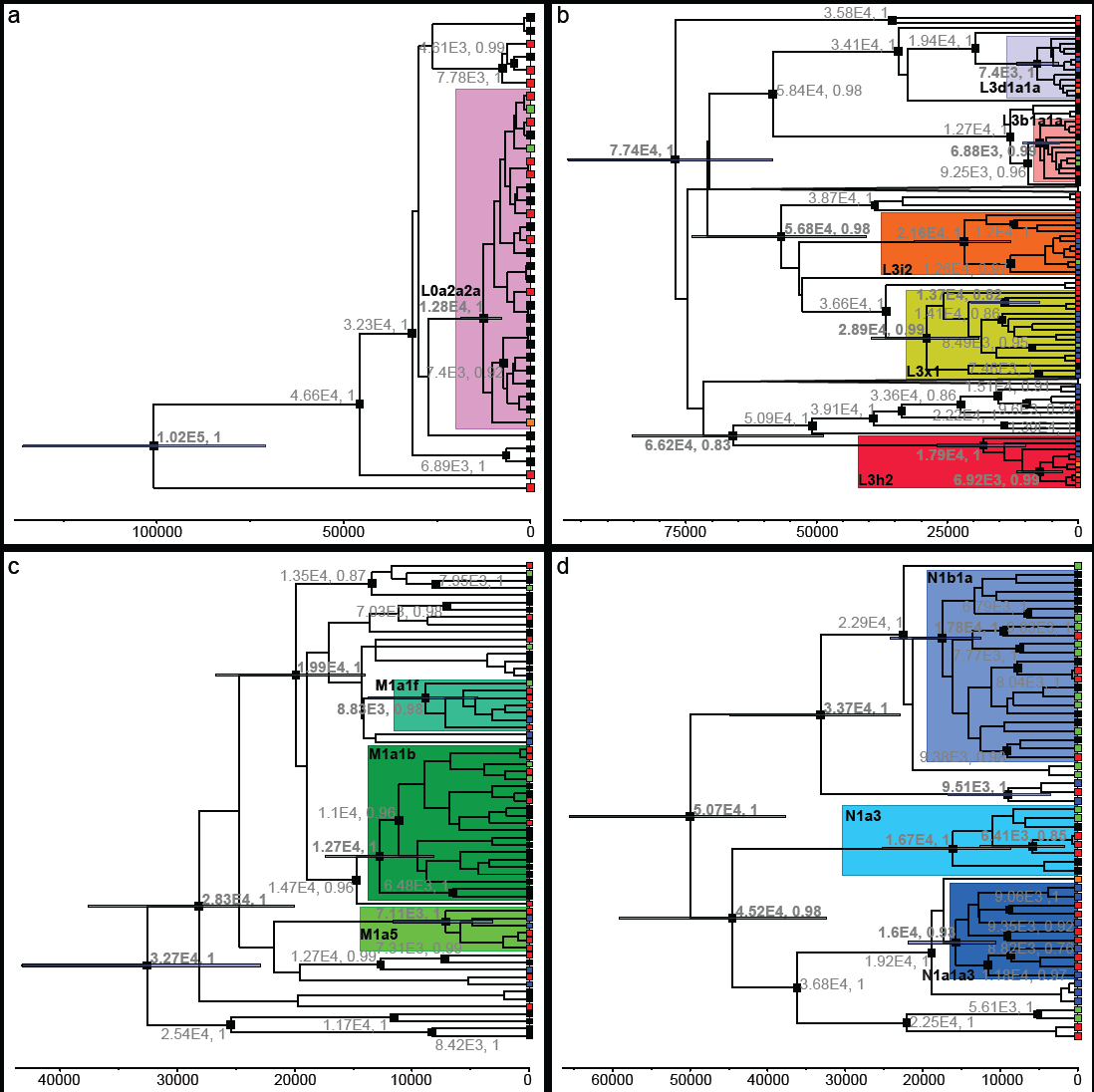
Detailed phylogenies of (a) L0a’b’f, (b) L3(xM,N), (c) M1, and (d) N1 are depicted. Subhaplogroups are colored as in Figure 1. Height 95% HPD bars are depicted for nodes listed in Table 1. Dates and posterior probabilities are provided for all nodes with ages older than 6 kya and posterior probabilities greater than 0.75. Tips are colored as follows: Red = Yemeni samples from the sample set of C.J.M.; Orange = Yemen samples from GenBank; Blue = East African samples (i.e., samples from the Horn of Africa and immediately neighboring countries); Green = samples from the Near East (i.e., samples from the Arabian Peninsula, the Levant, and Turkey); and Black = all other regions.

### Haplogroup L0

A primary goal in this study was to test Yemeni sequences for strict Yemeni monophyly, or more liberal Horn of Africa/southern Arabia monophyly, to investigate the possibility, and timing, of autochthonous evolution of ancient sequences in regions along the SDR. Six sequences (n=7 samples; divergence dates of identical sequences are estimated in BEAST and are conditional on the population size, substitution model, and molecular clock) were classified as subhaplogroup L0a2a2a. Bayesian phylogenetic analysis indicated that the Yemeni L0a2a2a sequences were not monophyletic; rather they were spread across the subhaplogroup and were interspersed with sequences of non-Yemeni origin including Arabia and Africa (Figure 2a). Additionally, most nodes in this group had low posterior probabilities limiting our ability to infer relationships within the group (Figure S2). For example, in some preliminary analyses, Yemeni sequences formed an Arabian clade with previously-published Saudi Arabian, Omani, or Yemeni sequences with low posterior probabilities that was not found in later analyses. The estimated divergence time for this subhaplogroup was 12.8 kya (95% HPD: 7.7-18.5 kya; Table 2). Given that Yemeni sequences in L0a2a2a group with both nearby and distant regions, this subhaplogroup is not a candidate for autochthonous evolution along the SDR and its divergence date is not sufficiently old to reflect the original migration out of Africa.

### Haplogroup L3(xM,N)

Four sequences (n=5) were classified as subhaplogroup L3b1a1a. Two sequences (n=3) formed a sister clade to the rest of L3b1a1a, while the remaining two were intermixed with sub-Saharan African and Near Eastern sequences (Figure 2b). Our phylogenetic analysis estimated the divergence date of this subhaplogroup to 6.9 kya (95% HPD: 3.6-10.5 kya; Table 1).

**Table 1.** List of key nodes from the complete phylogeny. Median ages, height 95% HPD intervals, and posterior probabilities of key nodes are listed and numbered as they appear in Figure 1. The twelve well-represented (n≥3) subhaplogroups analyzed in the current study have been shaded grey, and the quadrant of Figure 2 in which they are depicted has been indicated.

|  | **Node** | **Median Age (ybp)** | **95% HPD Interval (ybp)** | **Posterior Probability** |
| --- | --- | --- | --- | --- |
| 1 | Modern Human-Denisovan CA | 757,358 | (553,902 – 978,874) | 1.00 |
| 2 | Modern Human-Neanderthal CA | 389,303 | (287,858 – 499,301) | 1.00 |
| 3 | Neanderthal Common Ancestor | 121,547 | (94,916 – 149,202) | 1.00 |
| 4 | Maternal MRCA | 165,808 | (124,181 – 208,432) | 1.00 |
| 5 | L0a'b'f | 102,006 | (70,768 – 135,818) | 1.00 |
| 6 | L0a2a2a (Fig 2a) | 12,794 | (7,742 – 18,480) | 1.00 |
| 7 | L3 | 77,380 | (58,369 – 97,489) | 1.00 |
| 8 | M | 61,228 | (46,292 – 79,265) | 1.00 |
| 9 | N | 62,922 | (48,309 – 79,871) | 1.00 |
| 10 | R | 55,424 | (41,996 – 70,821) | 0.99 |
| 11 | L3e'l'k'x | 56,795 | (40,395 – 73,728) | 0.98 |
| 12 | L3b1a1a (Fig 2b) | 6,884 | (3,570 – 10,479) | 0.99 |
| 13 | L3d1a1a (Fig 2b) | 7,400 | (3,685 – 11,665) | 1.00 |
| 14 | L3h | 66,186 | (48,690 – 85,323) | 0.83 |
| 15 | L3h2 (Fig 2b) | 17,929 | (9,945 – 26,916) | 1.00 |
| 16 | *Yemeni/Somali clade within L3h2*<br>(Fig 2b) | 6,917 | (2,929 – 11,690) | 0.99 |
| 17 | L3i2 (Fig 2b) | 21,600 | (12,916 – 31,372) | 1.00 |
| 18 | L3x1 (Fig 2b) | 28,873 | (18,995 – 39,538) | 0.99 |
| 19 | *Yemeni clade within L3x1* (Fig<br>2b) | 13,693 | (7,335 – 20,548) | 0.82 |
| 20 | M1 | 32,740 | (22,953 – 43,254) | 1.00 |
| 21 | M1a | 28,313 | (19,995 – 37,636) | 1.00 |
| 22 | M1a1 | 19,940 | (14,032 – 26,709) | 1.00 |
| 23 | M1a1b (Fig 2c) | 12,666 | (8,166 – 17,434) | 1.00 |
| 24 | M1a1f (Fig 2c) | 8,828 | (4,403 – 13,715) | 0.98 |
| 25 | M1a5 (Fig 2c) | 7,106 | (3,126 – 11,631) | 1.00 |
| 26 | N1 | 50,733 | (37,640 – 65,557) | 1.00 |
| 27 | N1a | 45,192 | (32,419 – 59,191) | 0.98 |
| 28 | N1a1a3 (Fig 2d) | 16,001 | (10,513 – 21,895) | 0.93 |
| 29 | N1a3 (Fig 2d) | 16,697 | (8,636 – 25,159) | 1.00 |
| 30 | *Yemeni clade within N1a3* (Fig 2d) | 6,410 | (1,742 – 12,570) | 0.85 |
| 31 | N1b | 33,670 | (22,856 – 44,931) | 1.00 |
| 32 | N1b1a (Fig 2d) | 17,830 | (12,476 – 24,210) | 1.00 |
| 33 | N1b2 | 9,512 | (3,537 – 16,729) | 1.00 |

Four sequences (n=5) were classified as subhaplogroup L3d1a1a and these Yemeni sequences were found to be polyphyletic across the subhaplogroup and intermingled with sub-Saharan African sequences as well as a Pakistani sequence (Figure 2b). Additionally, many nodes in this subhaplogroup had posterior probabilities of <0.1, precluding any topological inferences (Figure S2). Our phylogenetic analysis produced a median estimate of 7.4 kya (95% HPD: 3.7-11.7 kya; Table 1) for the divergence of this subhaplogroup.

Four sequences were classified as subhaplogroup L3h2 (Figure 2b). Interestingly, these four Yemeni samples formed a clade with one Somali and two Yemeni samples gathered from GenBank (posterior probability = 0.99). Notably, two sequences previously published by Soares et al. (2012) (one Yemeni and one Somali) were identical to two of our sequences. Our phylogenetic analysis produced date estimates of the Yemeni/Somali clade of 7 kya (95% HPD: 2.9-11.7 kya; Table 1), while the subhaplogroup, as a whole, contained East African and Yemeni sequences and was estimated to date to 17.9 kya (95% HPD: 9.9-26.9 kya; Table 1).

Four sequences were classified as subhaplogroup L3i2, and these sequences were intermingled with Ethiopian, Somali and Omani sequences (Figure 2b). Our phylogenetic analysis produced an estimated divergence date for this subhaplogroup of 21.6 kya (95% HPD: 12.9-31.4 kya; Table 1).

A total of six sequences were classified as subhaplogroup L3x1, from which one was further classified as L3x1b. These sequences formed a clade with sequences from the Horn of Africa (Israeli sample EU092666 was identified by Behar et al. (2008) as an Ethiopian Jew) (Figure 2b). This subhaplogroup also contained a monophyletic clade of three Yemeni sequences dating to 13.7 kya (95% HPD: 7.3-20.5 kya; Table 1). The estimated divergence date for the whole subhaplogroup was dated to 28.9 kya (95% HPD: 19.0-39.5 kya; Table 1). Subhaplogroups L3h2, L3i2 and L3x1 are all candidates for autochthonous evolution along the SDR, although their divergence dates are not sufficiently old to reflect the original migration out of Africa.

### Haplogroup M1

A total of 20 sequences (n=21) were classified as haplogroup M1a and no sequences from sister-group M1b were found (Figure 2c). Thirteen sequences (n=14) were classified as M1a1 with four sequences further classified to M1a1b (n=5), and one sequence further classified to M1a1d. The phylogenetic analysis produced a median divergence date estimate for M1a1 of 19.9 kya (95% HPD: 14.0-26.7 kya; Table 1). Interestingly, five of the eight M1a1 sequences that did not belong to an established subhaplogroup formed a monophyletic group (posterior probability = 0.95) with one Ethiopian and one Saudi Arabian sequence with a median divergence date estimate of 8.8 kya (95% HPD: 4.4-13.7 kya, Table 1; Figure 2c). These seven mitogenome sequences possess a derived T6378C substitution not present in the other M1a1 sequences, and Pennarun et al. (2012) have suggested this lineage be named subhaplogroup M1a1f; we will henceforth use this notation for this clade and the samples within it. The remaining three M1a1 sequences were spread across the subhaplogroup (Figure 2c). Within M1a1b, Yemeni samples were intermixed with sequences from both nearby (e.g., Saudi Arabia) and distant regions (e.g., the Russian Caucasus) of western Eurasia (Figure 2c). Our phylogenetic analysis produced median divergence date estimates for the subhaplogroup of 12.7 kya (95% HPD: 8.2-17.4 kya; Table 1). M1a1b is not a candidate for autochthonous evolution along the SDR as its evolution is broadly geographically interspersed and its divergence dates additionally are not sufficiently old to reflect the original migration out of Africa.

Four sequences belonged to subhaplogroup M1a5 and the clade included sequences from Somalia and Ethiopia (Figure 2c). Our phylogenetic analysis produced a median estimate of 7.1 kya (95% HPD: 3.1-11.6 kya; Table 1) for the divergence of this subhaplogroup. Although this subhaplogroup’s divergence date is not sufficiently old to reflect the original migration out of Africa, it is a candidate for autochthonous evolution along the SDR. Notably, the final three M1a mitogenomes comprised a single M1a2 sequence and two M1a sequences belonging to no named subhaplogroups.

### Haplogroup N1

Fourteen sequences (n=16) classified as haplogroup N1 (including haplogroup I, which is nested within N1a1b), with 10 sequences (n=11) belonging to subhaplogroup N1a (including haplogroup I) and four sequences (n=5) belonging to subhaplogroup N1b. Amongst the N1a haplotypes, six sequences were classified subhaplogroup N1a1a3. Sequences from this subhaplogroup were intermingled with three Ethiopian and two Somalia samples (Figure 2d). This clade was estimated to date to 16 kya (95% HPD: 10.5-21.9 kya; Table 1). This subhaplogroup is a candidate for autochthonous evolution along the SDR but, as with the other groups mentioned above, its divergence date is too young to be related to the original migration out of Africa. Two Yemeni sequences (n=3) were classified as subhaplogroup N1a3 and formed a monophyletic group with a median divergence date of 6.4 kya (95% HPD: 1.7-12.6 kya; Table 1) and is closely related to sequences from both the Near East and Europe (Figure 2d). The remaining two N1a sequences belonged to haplogroup I, which is nested within N1a1b and sister to N1a1b1.

Within subhaplogroup N1b, three sequences (n=4) belonged to subhaplogroup N1b1a (of which one was further classified as N1b1a2) and were intermingled with other western Eurasian and North African sequences (Figure 2d). The N1b1a subhaplogroup had a median divergence date of 17.8 kya (95% HPD: 12.5-24.2 kya; Table 1). The remaining N1b sequence belonged to N1b2.

## DISCUSSION

We generated 113 mitogenomes from a diverse set of Yemeni samples and performed Bayesian phylogenetic analyses of these mitogenomes and 338 previously published mitogenome sequences to investigate the genetic relationship of Yemenis to neighboring populations. We were particularly interested in determining if AMHs may have migrated out of Africa along the SDR and if contemporary populations in Yemen, or in the greater region of Horn of Africa/southern Arabia, are descendants of these early migrants. We hypothesize that if early AMHs used the SDR, then contemporary descendant populations may carry basal, autochthonous lineages of the canonical OOA mtDNA haplogroups (i.e., L3, M, and N). We further hypothesize that autochthonous lineages indicative of the SDR would have diverged around 50-60 kya, when the original migrant populations swept across the landscape leaving descendant populations in their wake, after which mitogenome lineages carried by the descendant populations would have started to diverge from each other as they acquired population-specific substitutions.

In our study, we find the oldest monophyletic Yemeni clades, as well as the oldest monophyletic clades of Horn of Africa/Arabian sequences more broadly, in L3x1 with divergence dates of 13.7 kya and 28.9kya, respectively (Table 1). Both of these divergence dates are too young to reflect the initial migration out of Africa. Therefore, it is more likely that this lineage arrived in Yemen via a subsequent migration into the Arabian Peninsula, possibly via the SDR. The absence of ancient autochthonous lineages is good evidence that the contemporary Yemeni mitochondrial gene pool reflects more recent dispersal events than the expansion of AMHs out of Africa. The facts that ten of the twelve studied, well-represented (n≥3) subhaplogroups had median ages younger than 20 kya (Table 1) and that all but one of the Yemeni, or predominantly Yemeni, lineages (n≥3) within the subhaplogroups had median ages younger than 10 kya (Table 1, in grey) are consistent with recent (i.e., Holocene) migration playing a predominant role in shaping modern Yemeni mitogenomic diversity. Although other researchers, such as Thangaraj et al. (2005) and Macaulay et al. (2005), may have detected mitochondrial signals of ancient SDR migrations in regions such as the Andaman Islands or the Malay Peninsula, it is possible that recent migration into Yemen, in concert with genetic drift, has obscured any clear genetic signal for an early migration along the SDR. The signal of an SDR migration may have been left intact elsewhere due to various factors, such as geographic isolation limiting gene-flow or the stochastic nature of genetic drift on diversity.

Thus, our Yemeni data are relevant for discussions of more recent dispersals between the Arabian Peninsula, Africa, and other parts of Eurasia. Our results suggest that different types of migration have impacted modern Yemeni mitogenome diversity and we have grouped these dispersals into four types that are discussed below and are listed in Table 2. Briefly, the first type is recent migration into Yemen due to the Arab slave trade throughout sub-Saharan Africa. The second type of migration is recent (Holocene) East African migration and is similar to type 1, but differs in that Yemeni sequences cluster only with East Africans, while the third type of migration is older (Pleistocene) East African migration. The fourth type of migration is recent (Holocene) Western Eurasian migration characterized by subhaplogroups in which Yemeni sequences cluster with sequences from one or more regions of Western Eurasia.

**Table 2.**
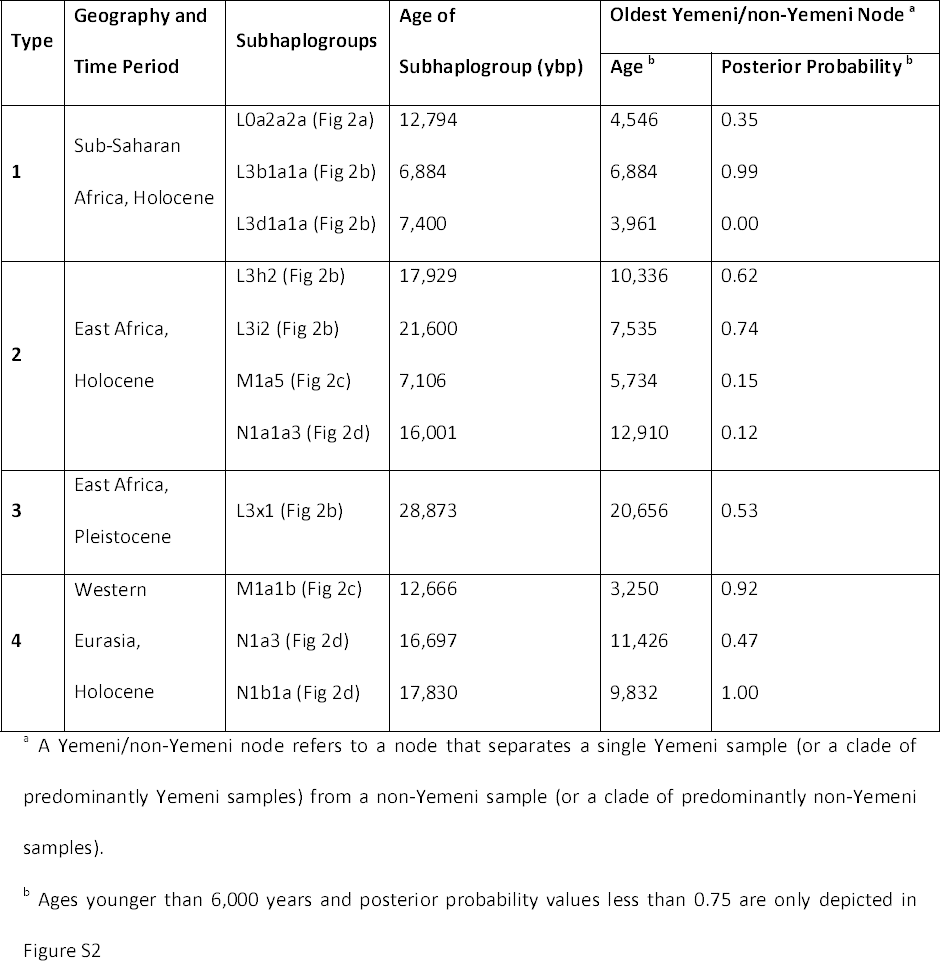
Four types of migrations inferred from our analyses. Representative subhaplogroups and associated ages and posterior probabilities are presented, and the quadrant of Figure 2 in which they are depicted has been indicated

The first type of migration likely reflects female-mediated migration from sub-Saharan Africa via the Arab slave trade over the last ˜2,500 years (Table 2). Richards et al. (2003) reasoned that the high frequency of African mitochondrial L lineages in Yemen is the product of female-mediated migration from sub-Saharan Africa via the Arab slave trade. This is discernible in our analyses of the African subhaplogroups L0a2a2a, L3b1a1a and L3d1a1a where Yemeni sequences clustered with sequences from multiple parts of sub-Saharan Africa, as well as from other populations with sub-Saharan African ancestry (e.g., African-Americans). During sample selection, we chose to sequence subhaplogroup L0a2a2a samples in addition to the L3(xM,N)/M/N samples because L0a2 was found at high frequency in our HVR I haplogroup data and we sought to determine if this lineage contained evidence of Yemeni monophyly, and thus *in situ* evolution in Yemen. However, our Yemeni L0a2a2a sequences did not cluster with each other and instead were closely related to other L0a2a2a sequences from many parts of Africa, which can be inferred as the effect of the slave trade. Additionally, we find two well-represented (n≥3) subhaplogroups from L3(xM,N) (L3b1a1a and L3d1a1a) that show a similar pattern of variation and may also indicate a recent dispersal of these subhaplogroups via the slave trade. From within these three subhaplogroups, we find that the median divergence dates for Yemen/non-Yemeni nodes [i.e., nodes separating a Yemeni or clade of (predominantly) Yemenis from a non-Yemeni or clade of (predominantly) non-Yemenis] are restricted to ˜5-7 kya. Considering that these are coalescent dates and would thus pre-date any actual migration to Yemen, these results are consistent with the arguments of Richards et al. (2003) that the Arab slave trade over the last ˜2,500 years has impacted modern Yemeni mitogenomic diversity.

The second type of migration that we found was evidence for recent East African migration in Yemen (Table 2). This is consistent with the multiple previous findings of recent migration events between East Africa and the Near East (Pagani et al. 2012; Boattini et al. 2013; Hodgson et al. 2014). We find this pattern in four of the twelve studied, well-represented (n≥3) subhaplogroups, of which two belong to L3(xM,N) [L3h2 and L3i2, which are said by Kivisild et al. (2004) to have originated in East Africa], one belongs to M1 (M1a5), and one to N1 (N1a1a3). In these subhaplogroups, our Yemeni sequences cluster almost exclusively with East African sequences and the oldest Yemen/non-Yemeni divergence dates in each of these groups range from 5-12 kya; the youngest of these four (M1a5) clearly stays within the Holocene (11.7 kya to present) while the oldest (N1a1a3) slightly traverses that boundary (Table 1: Type 2). It is important to note that many of these nodes within subhaplogroups have low posterior probabilities, and thus these dates should be interpreted with caution (Figure S2). Also, as macrohaplogroup L3 is thought to have evolved in Africa, the presence of 16 sequences from singleton/doubleton subhaplogroups from L3(× M,N) is strongly suggestive of additional recent African immigration into Yemen, beyond what we see in the well-represented (n≥3) subhaplogroups (Table 2, Types 1 & 2).

Only one subhaplogroup (L3 × 1) is supportive of our third type of older, East African migration (Table 2). Notably, this subhaplogroup is also thought to be East African in origin (Kivisild et al. 2004). The divergence dates for this subhaplogroup (Table 1) and the small Yemeni clade within it (n=3) (Table 1) are much older than all the other subhaplogroups we analyzed. In all of the other well-represented (n≥3) subhaplogroups, median divergence dates for Yemen/non-Yemeni nodes are restricted to the Holocene (11.7 kya to present), but our dates for L3 × 1 cross into the late Pleistocene. The age of our L3 × 1 subhaplogroup, as well as distribution of Yemeni sequences within it, may indicate back-and-forth migration between East Africa and Yemen during the Pleistocene, as opposed to more recent migrations.

Finally, in three subhaplogroups (M1a1b, N1a3, and N1b1a), Yemeni sequences are closely related to Near Eastern, European, and Caucasian sequences indicating the fourth type of migration (Table 2). In these groups, it seems both short-range and long-range Western Eurasian migration has occurred. Notably, within N1b1a, individual Yemeni sequences were closely related to sequences from all three regions. As is the case with migration types 1 and 2, Yemen/non-Yemeni nodes range from 3-10kya in these subhaplogroups. Additionally, in subhaplogroup M1a1f [as defined by Pennarun et al. (2012)] we found close relationships between Yemeni sequences and both a Saudi Arabian and an Ethiopian sequence, but we cannot infer the direction of migration (i.e., second vs. fourth), so we excluded it from Table 2.

### Eurasian Origin of Haplogroup M1

The presence of haplogroup M1a in Yemen is of particular interest as M1 is found at high frequency in East Africa and is thought to have either evolved in East Africa, or evolved in the Near East and been brought into Africa by back-migration (Olivieri et al. 2006). Olivieri et al. (2006) have argued for the latter explanation, and more specifically posit that M1 evolved in the Levant and was brought to Africa via migration, as opposed to being a marker of a back-migration along the SDR. Our findings are consistent with the explanation of Olivieri et al. (2006) as we found no M1b sequences in our Yemeni samples and if M1 was carried by an SDR back-migration through Yemen into Africa, then both M1a and M1b should be found in descendent populations along this route. In a SDR back-migration scenario, one would also expect Yemeni sequences to be basal to other M1a sequences, instead of being mixed within them as we found. These factors suggest that the presence of M1a in Yemen is the result of multiple immigration events, rather than a Yemeni evolution of M1a followed by emigration.

### Bayesian evolutionary methodology

Our study represents one of the first applications of Bayesian coalescent methods and relaxed molecular clocks for divergence dating (as implemented in BEAST) using whole mitogenome sequences, in contrast to most studies using contemporary mitochondrial samples that use only coding region data (e.g., Atkinson et al. 2008 and Gunnarsdottir et al. 2011). We performed multiple types of analyses to account for uncertainties in this new application of Bayesian methodology to partitioned human mitogenome data. We found that over-sampling from non-M1/N1 branches of macrohaplogroups M and N slightly widened our height 95% HPDs and suggest that this result may be related to the observation of Behar et al. (2012) that the molecular clock is being violated in these macrohaplogroups. Furthermore, the bases of these two groups are massive, unresolvable polytomies and the bifurcating structure of BEAST genealogies might inflate estimates above those made using other statistical methods (Soares et al. 2009). However, it could simply be that the relaxed clock model used in our analysis accurately reflects the substitution process in these macrohaplogroups and that our height 95% HPD intervals are wider because they accurately reflect the real uncertainty in date estimation, rather than being the product of model misspecification.

Importantly, over-sampling outgroup haplogroups (beyond what is needed for phylogenetic context) needs to be distinguished from over-sampling one’s haplogroups of interest, as the latter is necessary for proper phylogenetic analysis of these haplogroups and the former risks creating more uncertainty in estimates. Notably, our analyses did produce median date estimates largely consistent with previous work without using an undue number of calibrations. For example, we dated M1a to 28.3 kya (95% HPD: 20.0-37.6 kya; Table 1), which is similar to the coalescent estimate of 28.8 kya (±4.9 kya) by Olivieri et al. (2006). We also dated N1 to 50.7 kya (95% HPD: 37.6-65.6 kya; Table 1) which is similar to the ML estimate of 55,160 (95% confidence interval: 44,809-65,786) by Fernandes et al. (2012). Additionally, our median estimates for the origin of macrohaplogroups M and N are 61.2 kya (95% HPD: 46.3-79.2 kya; Table 1) and 62.9 kya (95% HPD: 48.3-79.9 kya; Table 1), respectively, which are consistent with other estimates that they arose around 60 kya (Atkinson et al. 2008; Behar et al. 2008; Soares et al. 2012). Significantly, the decrease in the width of node height intervals that we found when including two Denisovan mitogenomes suggests that additional independent calibrations, even in the form of serial samples from outgroup taxa, may provide significant information for calibrating substitution rates and improving the precision of divergence date estimates.

### Conclusions

We find mitochondrial evidence for multiple migration events into Yemen from sub-Saharan Africa, East Africa and western Eurasia that occurred relatively recently (i.e., primarily during the Holocene). While we did not find affirmative evidence for the SDR hypothesis, our data do not preclude or disprove the hypothesis as any genetic signal of this ancient event may have been overwritten by subsequent migrations (in both directions) or other demographic changes, as well as ongoing genetic drift. Hodgson et al. (2014) recently used simulation analyses to demonstrate that methods used to date admixture events are biased by recent admixture events (e.g., 300 or 900 years ago) relative to older admixture events (e.g., 1,500-6,000 years ago). This pattern of recent events erasing signatures of ancient events may have obscured evidence of ancient migration along the SDR. A test of this hypothesis would be to take a similar approach as implemented in our study to look for signatures of ancient migration in North African and Levantine mitogenomes as support for the NDR. If no support for migration along the SDR or the NDR can be found, it is likely that recent events have overwritten the genetic signature of ancient events since we know that AMHs did leave Africa, most likely through some part of the Arabian Peninsula whether or not a genetic signature of this event remains. Notably, nuclear genome sequence data from Yemen may prove informative for investigating the SDR. Ancient haplotypes may remain in modern Yemeni nuclear genomes, despite their absence from the mitogenome, in a similar manner to the preservation of introgressing Neanderthal haplotypes in modern non-African nuclear genomes but not in mitochondrial genomes (Green et al. 2008; Briggs et al. 2009; Green et al. 2010).

## MATERIALS AND METHODS

### Sample Mitogenome Sequencing

For this study, 113 Yemeni mitogenomes from the collection of C.J.M. were sequenced (GenBank accession KM986515 – KM986627). Using previously generated HVR I sequences, a basic maximum likelihood (ML) tree of the 550 samples in the collection was generated using PhyML (Guindon et al. 2010), from which an initial set of 23 diverse sequences representative of the breadth of mitogenomic diversity in Yemen was chosen for whole mitogenome sequencing. Mitogenomes from these 23 individuals were amplified per Gonder et al. (2007) and sequenced by means of Sanger sequencing at the University of Florida’s Interdisciplinary Center for Biotechnology Research (ICBR). Sequences were assembled with the revised Cambridge Reference Sequence (rCRS) (Andrews et al. 1999) using Sequencher v5.0 (http://genecodes.com/).

An additional set of 90 mitogenomes was generated using indexed, pooled library construction and Illumina HiSeq 2000 sequencing. This set focused on samples classified to haplogroups L3(xM,N) (n=39), M (n=29), N (n=13) and L0a2 (n=9) based on HaploGrep classification (Kloss-Brandstätter et al. 2011). All 90 Yemeni samples (as well as four contamination control samples) were sheared to 200bp fragments using the Covaris DNA shearing system E210 (http://covarisinc.com) following the manufacturer’s specifications. Following shearing, DNA was purified using SPRI beads, which we generated in our lab following the protocol set forth by Rohland and Reich (2012). Sheared samples underwent indexed library preparations, using an approximately 200bp PCR product for the positive control, following Meyer and Kircher (2010). All libraries were pooled in equimolar ratios with 200 ng of DNA per library. Per Meyer et al. (2007) and Maricic et al. (2010), mitochondrial bait was generated using two long-range PCR amplifications using two primer pairs designed to yield minimum overlap; the bait was amplified from a salivary DNA sample not used for mitochondrial sequencing. The long-range PCR products were purified, quantified, equimolarly pooled, and sheared using the Covaris E210 to 400 bp following the manufacturer’s specifications. Biotinylated adapters were ligated to the bait and then bonded to streptavidin beads. The pooled, indexed libraries were hybridized with the bait at 65°C for 48 hours, as in Maricic et al. (2010). Enriched libraries were eluted, quantified using qPCR, and amplified and purified. These enriched libraries were sequenced at the Illumina core lab at the University of Miami, Miller School of Medicine on an Illumina HiSeq 2000 with 100bp pair-end read sequencing. A total of 2 µg of library DNA was used for sequencing. Sequence data were separated by index by the sequencing lab. Following index separation, the program FastQC (available at http://www.bioinformatics.babraham.ac.uk/projects/fastqc/) was used to assess the quality of each fastq file. Adapter sequences and primer sequences were trimmed from reads using fastq-mcf, and paired-end reads were joined using the program fastq-join (http://code.google.com/p/ea-utils).

For each sample, the program Bowtie 2 (Langmead and Salzberg 2012) was used to align reads to the rCRS; reads that did not align to the rCRS were judged as non-mitochondrial in origin and excluded from assembly. Using only the reads aligning to the rCRS, mitogenomes were assembled using the program Mapping Iterative Assembler (MIA) previously described by Briggs et al. (2009). All modern human mitogenomes (including comparative mitogenomes described below) were assigned to haplogroups according to PhyloTree mtDNA tree Build 15 (rCRS version) using MitoTool (van Oven and Kayser 2009; Fan and Yao 2013). In cases where more than one haplogroup could be assigned, PhyloTree was consulted on a case by case basis. In all other cases, the MitoTool haplogroup assignment was accepted. All samples (including comparative mitogenomes; see below) were aligned to each other for further analysis using MAFFT v7.050 (Katoh and Standley 2013) and curated by hand to ensure proper alignment.

### Bayesian Phylogenetic Analysis

The set of 113 mitogenomes were subjected to Bayesian evolutionary analysis using BEAST v1.7.2 (Drummond et al. 2012). Additional, previously published Yemeni mitogenomes from the collection of C.J.M. are included as well: 9 were in included in all analyses, and another 12 were included in the analyses with additional non-M1/N1 samples (further described below) (Musilová et al. 2011; Černý et al. 2011; Al-Abri et al. 2012) (see Table S2 for all Yemeni samples). A set of 215 previously published, comparative modern human mitogenomes (nearly all non-Yemeni), as well seven Neanderthal and Denisovan mitogenomes, were included in the Bayesian analyses and an additional set of 95 samples were also included in analyses with additional non-M1/N1 samples (further described below) (see Table S3 for all modern comparative samples). Comparative samples were specifically chosen from subhaplogroups for which we have three or more Yemeni mitogenomes, as well as nearest sister subhaplogroups, in order to rigorously test the monophyly of Yemeni mitochondrial haplogroups, estimate the age of the subhaplogroup, and estimate the date of any monophyletic Yemeni clade(s) within it. Sequences from other major haplogroups within macrohaplogroups M and N (e.g., CZ, M29’Q, R) were included to help define the roots of macrohaplogroups M and N. Five Neanderthal mitogenomes from dated remains from Vindija (NC_011137), Feldhofer (FM865407, FM865408), El Sidron (FM865409), and Mezmaiskaya (FM865411) were included in all analyses as an outgroup clade (Green et al. 2008; Briggs et al. 2009). In some analyses, two Denisovan mitogenomes (FN673705, FR695060) were included to form an additional outgroup (Krause et al. 2010; Reich et al. 2010).

Phylogenetic analyses were run using BEAST v1.7.2 (Drummond et al. 2012) on the University of Florida High Performance Computing Center’s HiPerGator cluster. A coalescent tree prior with constant population size was used in our analyses (Kingman 1982; Drummond et al. 2002). The coding and control regions were partitioned for these analyses. The General Time Reversible (GTR) model (Tavaré 1986) with invariant sites and gamma distributed rate variation (GTR+I+G) was used for the coding region partition, while the Tamura and Nei (1993) (TN93) model with invariant sites and gamma distributed rate variation (TN93+I+G) was used for the control region partition (initial analyses with the GTR model applied to the control region partition failed to converge). A normal distribution with a mean of 1.691×10^-8^ substitutions per site per year (Atkinson et al. 2008) and a standard deviation of 2.5×10^-9^ substitutions per site per year was used as the prior probability distribution for the substitution rate for the entire mitogenome while a scaleless, uninformative prior was placed on the mean of the uncorrelated lognormally-distributed relaxed clock model (Drummond et al. 2006) from which branch rates were drawn. Additionally, a prior distribution was set on the base of the modern human clade (i.e., maternal MRCA), which had a normal distribution centered around 150,000 years with a standard deviation of 50,000 years, to provide additional calibration and better inform the molecular clock. Based on the topologies of preliminary analyses, some haplogroups were constrained to be monophyletic (van Oven and Kayser 2009). Analyses were run for 200,000,000 generations, sampled every 10,000 generations, and replicated at least once, while all log files were analyzed in Tracer v1.5 (http://tree.bio.ed.ac.uk/software/tracer/) to ensure Markov chain convergence. All tree files were analyzed using TreeAnnotator v1.7.2 with settings to use median node heights with a posterior probability limit of 0.5 and to discard the first 40,000,000 generations (20%) as burn-in. Consensus trees (i.e., maximum clade credibility trees) were visualized using FigTree v1.4.2 (http://tree.bio.ed.ac.uk/software/figtree/).

Considering that the main set of 342 or 344 (with Denisovan) samples was biased towards M1 and N1 (relative to other branches of macrohaplogroups M and N), analyses were also performed with 107 additional samples, specifically chosen to not belong to M1 or N1 but instead to represent other branches of M and N (Tables S1, S2), in order to test whether variation in sample sizes of sequences from different branches affected node heights. Additionally analyses were also performed with data sets that included and excluded ancient Denisovan mitogenome sequences in order to assess the effect of ancient and distantly related, serially sampled sequences on the calibration of evolutionary rates and the estimation of ingroup divergence times. In order to test whether differences between analyses with and without Denisovans were due to the Denisovan sequences or simply due to having two extra samples included, 10 replicates of the analysis that included the Denisovan sequences (n=344 samples) were run, but each with a random pair of non-Yemeni modern human samples removed (thus n=342). Dating results from these replicates were then compared to the original Denisovan-including analysis (n=344 samples) and the corresponding Denisovan-lacking analysis (n=342). As a result of these two sets of comparisons, analyses were performed in quadruplicate on four subsets of the total mitogenome sequence dataset. Half of the analyses contained the additional samples from other branches of M and N and half of the analyses included Denisovan mitogenome sequences, yielding four distinct, overlapping data sets.

In order to complete these comparisons, all maximum clade credibility tree files were first loaded in R v.3.0.1 (R Core Team 2014) using the ape (Paradis et al. 2004) and Phyloch (http://www.christophheibl.de/Rpackages.html) packages, so that all node information could be extracted. For comparisons of analyses with and without additional samples from non-M1/N1 branches, we compared mean width of all the height 95% HPD intervals (Table S4). For comparisons of analyses with and without Denisovans, node heights and the corresponding 95% highest probability density (95% HPD) intervals for 14 major nodes of varying ages were compared, as well as the mean width of all the height 95% HPD intervals on each tree. For these 14 nodes, it was noted whether the Denisovan-lacking analysis had an older or younger node height and narrower or wider height 95% HPD intervals than the 10 replicate analyses. It is also important to note that, in all cases where the mean width of all the height 95% HPD intervals on the trees was calculated, the node separating Denisovans from modern humans and Neanderthals (as well as the daughter node separating the two Denisovans) was excluded from these calculations. This was done as the node separating the Denisovans from modern humans and Neanderthals is the oldest divergence on the tree and has the widest height 95% HPD interval and, thus, greatly biases the overall mean height 95% HPD interval width to be wider when compared to analyses without Denisovans (which lack these nodes).

We found that the analyses including additional samples from non-M1/N1 branches of macrohaplogroups M and N yielded comparable dating results to analyses lacking these extra sequences, indicating that our largely M1/N1 data sets did not bias our divergence date estimates although it was noticed that the datasets with the additional sequences generated slightly wider 95% HPD intervals (Table S4). Analyses of data sets with Denisovan mitogenomes (n=344 and n=451) produced phylogenies where many divergence dates on the trees decreased very slightly (i.e., the median node heights were more recent) and the mean width of all the height 95% HPD intervals were narrower when compared to otherwise identical analyses that lacked Denisovan sequences (n=342 and n=449) (Tables S3, S4). Also, when we compared replicates of the Denisovan-including analysis with random pairs of modern human sequences removed (thus n=342) to the original Denisovan-lacking analysis (n=342) (thus ensuring equal sample sizes) at the 14 studied nodes, all the Denisovan-including analyses had younger dates (as noted by median node heights) and narrower height 95% HPD intervals than the Denisovan-lacking analysis (n=342) (Table S5). Additionally, the mean widths of all the height 95% HPD intervals were still consistently narrower when Denisovan sequences were included (Table S5).

All subsequent results reported in this study are from the analysis that excluded the additional non-M1/N1 sequences and included the Denisovan sequences (n=344). Additionally, this analysis was re-run in TreeAnnotator v1.7.2 with a posterior probability limit of 0.0 (instead of 0.5), so that node heights for low posterior nodes of interest could be calculated (e.g., Yemeni/non-Yemeni divergences within subhaplogroups).

## ACKNOWLEDGMENTS

Appreciation goes to the Yemeni people for their participation in this study. Thanks also go to the University of Florida High Performance Computing Center and the HiPerGator cluster support staff for their assistance in assembling sequences and conducting analyses. Mitogenome sequences generated in this study have been submitted to GenBank (http://www.ncbi.nlm.nih.gov/genbank) with accession numbers KM986515 – KM986627. This work was supported by the National Science Foundation (BCS 0518530 to C.J.M. and BCS 1258965 to C.J.M and D.N.V.) and by a Howard Hughes Medical Institute Distinguished Mentor Award to C.J.M. The authors have no conflicts of interest.

## SUPPLEMENTARY TABLE/FIGURE LEGENDS

**Table S1.** List of 90 Yemeni samples sequenced using Illumina HiSeq 2000 with number of reads (after joining paired ends), the percent of reads aligning to the rCRS using Bowtie2, and the mean, minimum, and maximum sequence coverage for each of the sample

**Table S2.** List of Yemeni samples analyzed, with haplogroup classification, GenBank accession number, Yemeni governate, and sequencing method listed

**Table S3.** List of comparative modern human mitogenome sequences analyzed, with haplogroup classification, GenBank accession number, country/region, and source listed

**Table S4.** Comparisons of phylogenetic analyses of different datasets. The effects of including/excluding additional non-M1/N1 and including/excluding Denisovan samples on the mean width of all height 95% HPD intervals are presented.

**Table S5.** Analyses of the effects of including Denisovan sequences on tree node height and height 95% HPD interval widths for tree-wide means as well as for 14 major nodes of varying ages

**Figure S1.** Results on 90 sequenced Yemeni samples depicting the number of reads that do and do not align to the rCRS

**Figure S2.** Entire phylogeny with dates and posterior probabilities for all nodes is depicted. All branches of macrohaplogroup M are colored dark green and all branches of macrohaplogroup N are colored dark blue. Subhaplogroups are colored as in Figures 1 and 2. Tip labels have been colored as such: Red = Yemeni samples from the sample set of C.J.M.; Orange = Yemen samples from GenBank; Blue = East African samples (i.e., samples from the Horn of Africa and immediately neighboring countries); Green = samples from the Near East (i.e., samples from the Arabian Peninsula, the Levant, and Turkey); and Black = all other regions.

**Figure S3.** Entire phylogeny with height 95% HPD bars for all nodes is depicted. All branches of macrohaplogroup M are colored dark green and all branches of macrohaplogroup N are colored dark blue. Subhaplogroups are colored as in Figures 1 and 2. Tip labels have been colored as such: Red = Yemeni samples from the sample set of C.J.M.; Orange = Yemen samples from GenBank; Blue = East African samples (i.e., samples from the Horn of Africa and immediately neighboring countries); Green = samples from the Near East (i.e., samples from the Arabian Peninsula, the Levant, and Turkey); and Black = all other regions.

